# Paving the way towards understanding the inflammatory pathways triggered by giant viruses in mammalian cells: effect of mimivirus-cell interactions on IκBα degradation

**DOI:** 10.1101/2021.09.16.460633

**Authors:** Juliana dos Santos Oliveira, Dahienne Ferreira Oliveira, Victor Alejandro Essus, Gabriel Henrique Pereira Nunes, Leandro Honorato, Leandro Oliveira, Leonardo Nimrichter, José Mauro Peralta, Allan Jefferson Guimaraes, Debora Foguel, Juliana Reis Cortines

## Abstract

Even after two decades since the identification of the first giant virus, the Acanthamoeba polyphaga mimivirus (APMV), it still elude scientists. Their gigantic size and genome are unique in the whole virosphere, and many aspects of their biology are still unknown, including their possible hosts. They are cultivated in laboratories using *Acanthamoeba* cells as hosts, but little is known about the infectivity of these giant viruses in vertebrate cells. However, there is evidence of the possible involvement of APMV in pneumonia and activation of inflammatory pathways. Among the hundreds of prospected giant viruses members is Tupanvirus, isolated in Brazil. Its particles have a characteristically large size varying between 1.2 to 2 μm and are covered by fibrils. In the present work, we aim to study the consequences of the incubation of APMV and Tupanvirus with mammalian cells. These cells express Toll-like receptors (TLR) that are capable of recognizing lipopolysaccharides, favoring the internalization of the antigen and activation of the inflammatory system. We used a lineage of human lung adenocarcinoma cells (A549) to evaluate possible effects of TLR activation by the giant viruses and if we could detect the probable cause of the said giant-virus dependent pneumonia. Our results show that APMV and Tupanvirus (TPV) activate cellular receptors related to the Toll-like 4 type-induced inflammatory response and that the A549 cells are capable of internalizing the latter virus. Therefore, this study brings new insights into the possible interactions established between mimiviruses (here represented by APMV and Tupanvirus) and members of the innate cellular immune response.

## 1 Introduction

The Acanthamoeba polyphaga mimivirus (APMV) was first isolated in 1992 during a pneumonia outbreak in the city of Bradford, United Kingdom. However, its identification and classification as a new type of virus would only occur in 2003, with its discovery heralding a new era in virology. Among its many distinguishing features, the APMV possesses a pseudo-icosahedral capsid with a size nearing 500 nm in diameter. Its particle is almost entirely covered by fibrils (125 - 140 nm), save for the stargate, feature that plays a role in the liberation of the virion internal contents, characterized by the presence of a star-shaped seal-like structure (La Scola et al., 2003; Schard et al., 2020; dos Santos Oliveira et al., 2021). Besides the gigantic particle size, its genome is also unique, being formed by a double-stranded DNA nearing 1.2 Mpb in size and encoding genes related to the expression of over 1000 proteins, transporter RNA (tRNA), tRNA aminoacylation, and non-coding RNA, all never seen before in sequenced viral genomes (La Scola et al., 2003; Raoult et al., 2004, Legendre et al., 2011). Considered the first giant virus (GV), its morphological and genetic characteristics are so peculiar that they impeded the phylogenetic insertion of the APMV in a previously established taxonomic family, making necessary the creation of the new *Mimiviridae* family and the *mimivirus* genus (Aherfi et al., 2016; Colson et al., 2017a). The isolation of APMV was a hallmark discovery in the field of virology and triggered a worldwide search for giant viruses culminating in the discovery of hundreds of new species. Among these discoveries are the Tupanvirus (TPV), which were isolated in 2018 in Brazil and are members of the *Mimiviridae* family, with currently only two known strains (TPV-soda lake and TPV-deep ocean) that compose the *tupanvirus* genus. Similar to the APMV, it possesses a capsid with pseudo-icosahedral geometry near 450 nm in diameter, almost totally covered in fibrils, and presents the stargate in one of its vertices. It also possesses a large dsDNA genome (∼1.5 Mpb) that encodes near 1.276 proteins, aminoacyl tRNA synthetases, tRNA, factors associated with tRNA/mRNA translation, and maturation (Abrahão et al., 2018). However, a unique aspect of TPV that distinguishes it from other giant viruses: the presence of a long cylindrical tail at the opposite side from the stargate vertice that is also covered by fibrils and portraits variable lengths (Abrahão et al., 2018).

Although their function is still unknown, it is widely believed that fibrils play an important role in GVs infection cycles. They are found in almost all the different (known) GV families (with exception of Pacmanviruses, Cedratviruse*s*, and Pithoviruses), although they are located at distinct sites in the particle (dos Santos Oliveira et al., 2021). Data published by Kuznetsov and collaborators have shown through atomic force microscopy that the APMV capsid surface is coated by a protein matrix with high levels of sugar. Peptidoglycan protrusions might act as anchoring points for the fibrils and in their protection (Kuznetsov et al., 2010, 2013). Although the characterization of the fibril matrix has yet to be completed, the existence of peptidoglycans is sustained by the APMV susceptibility to Gram coloration, and due to the presence of the *L136* gene (Piacente et al., 2012, 2017). This gene encodes an enzyme that acts in the synthesis of viosamin, a monosaccharide found in the O-antigen subunit of the lipopolysaccharide (LPS) of many bacterial strains. It was also evidenced by the presence of rhamnose in the fibrils, another monosaccharide present in bacterial LPS (Parakkottil Chothi et al., 2010; Piacente et al., 2012). The fibrils are disposed in a dense layer along most of the virion surface, in the case of the members of *Mimiviridae* family. They seem to act in the virus-cell contact, which makes them more attractive for professional phagocytes, such as amoebas (Rodrigues et al., 2015).

The large size of these viruses (> 500 nm) has led Ghigo and collaborators (2008) to propose, after an ample study, that in the vast majority, their replicative cycles were dependent on the virus-cell entry through phagocytosis (Ghigo et al., 2008). In fact, professional phagocytic cells such as certain amoebal trophozoites with intense phagocytic activity are the main GVs hosts used in laboratory (Colson et al., 2017b; Oliveira et al., 2019a; Yaakov et al., 2019). However, the literature has yet to enlighten the intricacies of GV internalization by amoebas. It’s thought that certain portions of the capsid or fibrils of APMV or TPV can activate cellular receptors that trigger cytoskeleton changes, allowing for the phagocytosis of the virions (Ghigo et al., 2008; Ostrowski et al., 2016). In humans, the phagocytic process can be undertaken by professional (i.e. macrophage), non-professional (i.e. epithelial cells) and specialized (i.e. Sertoli testicle cells) phagocytic cells (Arandjelovic and Ravichandran, 2015). The similar structural aspects between the fibrils and bacterial structures can be related to the host cellular response, since LPS, bacterial membrane structures and viral components interact in interesting ways with receptors from the Toll-like (TLRs) family (Vaure and Liu, 2014).

The TLR family of receptors is responsible for the identification of pathogen-associated molecular patterns (PAMP), being present in the front line of the innate immune system response (Vaure and Liu, 2014). Currently, more than 10 types of TLR in mammals are described with significant homology to human TLRs. The TLR4 are proteins composed of intra- and extracellular domains located in the membrane of cells from the immune system, central nervous system, endothelium, myocardium, and pulmonary alveolar epithelium (O’Neill et al., 2013). These are sensitive to the presence of LPS, and when activated can activate two signaling pathways: one dependent on the primary response protein of the myeloid 88 differentiation (MyD88) and the MyD88 independent pathway. In the first pathway, the IRAK4 kinase (interleukin-1 receptor-associated Kinase 4) suffers phosphorylation and activates IRAK1 (interleukin-1 receptor-associated Kinase 1). The complex formed by MyD88 and the interleukin 1 and 4 recruit TRAF6 (TNF receptor-associated factor 6) so that the complex TAK1 (TGFβ activated kinase 1) and TAB2 and 3 (TAK1 binding protein) can activate the kinases IKKα, IKKβ and IKKγ. These three kinases promote the IκB phosphorylation and its degradation allows the translocation of the nuclear factor-κB (NF-κB) to the nucleus, and the production of inflammatory cytokine. The MyD88 independent pathway needs adapter molecules TRIF (TIR domain containing adapter protein inducing interferon IFN-β) and TRAM (TRIF-related adaptor molecule) to activate a distinct set of transcription factors (O’Neill et al., 2013; Baizabal-Aguirre et al., 2016). Based on the role these types of receptors have in the cellular immune response and the identification of antigens such as LPS, it is possible that it is involved in the cellular response to GVs.

Although there is still a need for more conclusive evidence, previously published data suggest that APMV might be able to cause viral pneumonia (Khan et al., 2007; Saadi et al., 2013; Zhang et al., 2016). Serological tests and western blot assays showed the presence of specific antibodies and viral proteins for APMV and other members of the *Mimiviridae* family in samples collected from patients that were afflicted with pneumonia. As such, it is paramount that the possible impacts these viruses can have in humans be extensively explored. In this context, the present work evaluates the propensity of the APMV and TPV-soda lake internalization by adenocarcinomic human alveolar basal epithelial cells (A549). This cell line was chosen due to serving as an excellent model of alveolar epithelium type II, which may shed light on the induction of pneumonia by GVs. We also evaluated changes in TLR4 expression and IkB degradation in cells kept in the presence and/or absence of different multiplicity of infection of APMV and TPV. Since APMV and TPV fibrils can be composed of protein content similar to LPS, it is suggested that short incubation periods can be enough to cause a response from both of these indicators, resulting in an important inflammatory process.

## 2 Materials and Methods

### Viral particles’ purification

Three million *Acanthamoeba castellanii* cells (strain 30011) were grown in 75 cm^2^ cell-culture flasks and maintained in Peptone, Yeast Extract and Glucose medium (ATCC Medium 712) containing 1 mg/ml Penicillin/Streptomycin (Sigma-Aldrich, USA) and 15 μg/ml Gentamicin (Thermo Fisher Scientific, USA) for one hour. After complete adherence, the amoebas were infected with APMV or TPV-soda lake (TPV-SL was used for all experiments in the present work) at a multiplicity of infection (MOI) of 0.5 and stored in an oven at 28ºC. After 96 hours of infection, the collected cell lysate was carefully placed in 22% sucrose cushion, and centrifuged at 36,000 ×g for 30 min. The resulting viral pellets were resuspended in 500 μL of sterile PBS. Virus titers were determined using the method of Reed and Muench (Reed and Muench, 1938). On average, purification of APMV yielded titers of 10^8^ TCID50/ml and TPV of 10^7^ TCID50/ml.

### Labeling of viral particles

APMV and TPV-SL stocks at the desired MOI were centrifuged (25,000 *xg*, 10 min, 4°C) and the viral pellet was separated. Rhodamine-phalloidin (25 μg/ml) diluted in 1 ml of sterile PBS was filtered in a 0.22 μm filter and added to the desired virus pellet. The pellet is then resuspended in this marking solution and submitted to orbital shaking (Proenix Tecnologias) for 1 h at room temperature. Unbound rhodamine was removed by 5 washes with 1 ml of sterile PBS, after centrifugation under the conditions cited above. The resulting pellet was resuspended in PBS in the original stock volume.

### Cell culture

The A549 cell line (ATCC CCL-185) derived from human lung adenocarcinoma cells was chosen to carry out this work. Cells were grown in 75 cm^2^ cell culture flasks, maintained in Dulbecco’s Modified Eagle’s Medium (DMEM - Sigma Aldrich) supplemented with 10% fetal bovine serum plus 1% Gentamicin and stored in an incubator containing 5% CO_2_ at 37°C. Culture medium replacements were performed 2-3 times per week. In experiments involving polymyxin B (SIGMA, USA), cell cultures were previously treated with 50 μg/ml of this compound for one hour. Soon after this incubation, A549 cells were immediately exposed to DMEM, LPS (100 ng/ml), APMV (MOI 50) or TPV (MOI of 10 or 50).

### Confocal Fluorescence Microscopy

Ten thousand A549 cells were plated in 24-well plates. After 24 hours, the cells were incubated with TPV (MOI of 10 or 50) and APMV (MOI of 50) for 6 hours and kept in DMEM for another 36 hours. Then, the cells were fixed on coverslips present in 24-well plates, with 4% paraformaldehyde for 15 minutes at room temperature. Cells were blocked with 5% BSA, in PBS pH 7.4 for 1 hour, followed by the incubation with fluorescein isothiocyanate (FITC)-cholera toxin B subunit (CTB; Sigma, C1655, 1: 1000 diluted in sterile PBS) for 1 hour at 4°C, washed 3x with PBS and then incubated with DAPI (1:5000) for 15 minutes, washed again 3x with PBS and mounted with Prolong® on slides for microscopy analyses. Images were acquired with a Leica TCS-SPE confocal microscope equipped with an oil immersion objective 63× and a Nikon DS-fi2 camera operated with standard QC capture software (Leica, USA). Image analysis was done using the ImageJ program.

### Protein expression analysis by Western blotting

The expression of TLR4 and IkBα proteins were analyzed using Western blotting assays. Cultures of A549 cells, containing 10^6^/well in six-well plates, were incubated with APMV (MOI 50) and TPV (MOI 10 and 50) and LPS (100 ng/ml) at times of 30 minutes. Polyacrylamide gel electrophoresis (PAGE) with sodium dodecyl sulfate (SDS) was loaded with 150 μg of total protein from cell media. The proteins were separated by electrophoresis and transferred to polyvinylidene fluoride membranes (Millipore). The membranes were incubated with appropriate mouse antibodies directed against TLR4 (ab22048, 1:1000 diluted in Odyssey buffer, abcam) e IκBα (Cat #: 9247, 1:1000 diluted in Odyssey buffer, Cell Signaling Technology). After incubation with the respective secondary mouse antibodies (IRDye,680 Licor, USA) immunoreactive signals were detected by Fluorescence. Fluorescence was measured with odyssey FC Imaging System (Licor). Densitometry was normalized using GAPDH as a loading control. Statistical analysis was done using the Student T test calculated. The data was evaluated as a ratio of the target proteins and the relative expression of the control protein. Statistical significance was determined when p ≤0,05.

### Cell viability assay

The Live / Dead® Qualitative Fluorescence Assay Kit (Molecular Probes) was used to qualify the detection of A549 viability. A549 cell cultures in 96-well plates were incubated with TPV (MOI 0.1, 1, 10, 50) for 6 hours. The viral solution was removed and cultures were maintained with DMEM for 36 hours. After the incubation time, the cells were washed 3X with PBS and incubated with Hoechst (1: 5000, nuclear dye) for 15 min, then the cells were washed 3X PBS and incubated with Calcein AM (1: 1000, marker of living cells) and Ethidium homodimer-1 (EthD-1) (1:1000, dead cell marker) and observed by reverse fluorescence microscopy ImageXpress micro-XL (Molecular Advices) with a specific program (MetaXpress 6.0, Molecular Advices). Quantitative analyzes were made from the ratio of cells plated to dead cells.

## 3 Results

### A549 cells interact with APMV and TPV

A549 cells are excellent models for studies of respiratory infections, as they occasionally play the role of replication platforms for the viral pathogens that cause these infections. In this scenario, we chose to study the interactions that could be established between GVs and A549 cells. Through fluorescence microscopy, we analyzed the ability of A549 cells to interact with APMV and TPV. Thus, we kept the cell cultures in contact with DMEM medium containing APMV (MOI 50) or TPV (MOI 10 and 50) labeled with rhodamine for six hours, then replaced by DMEM supplemented with bovine serum and antibiotics, for another 36 hours. The resulting microscopy images are shown in **Figure 1** (and in **Supplementary Material**) and suggest that the particles of the two viruses are internalized by this cell line. **Figure 1A** shows the control group of cells that were not exposed to the viruses and were kept in DMEM only. **Figure 1B** represents the cells exposed to APMV in an MOI of 50, where it can be observed that a great quantity of viral particles appear to be co-localized with the cell membrane. The same was observed in **Figures 1C** and **1D**, that represent the cells exposed to TPV in MOI of 10 and 50, respectively. The number of viral particles along the cells increased proportionally to the increase in MOI used, as can be seen by the increasing number of red dots in the microscopies. For simplicity, we considered each dot to be a presumed particle number. When counted, there were a total of 352 dots in the samples exposed to APMV at MOI of 50 (**Figure 1B**), 34 and 447 dots in samples exposed to TPV at MOI 10 (**Figure 1C**) and 50 (**Figure 1D**), respectively. The increase of over 10x in the numbers of dots between the MOIs used in the TPV sustains the observation of the proportional increase between MOI and virus entry. We also observed overlapping points of viruses with the nucleus, but it can’t be confirmed if there was an invasion of such organelle. The co-localization of both viruses on the cell membrane is a strong indicator that the cell-virus contact occurred. Also suggests that the fibrils are recognized by cell receptors, which would make A549 cell lines susceptible to GVs.

**Figure 1:**
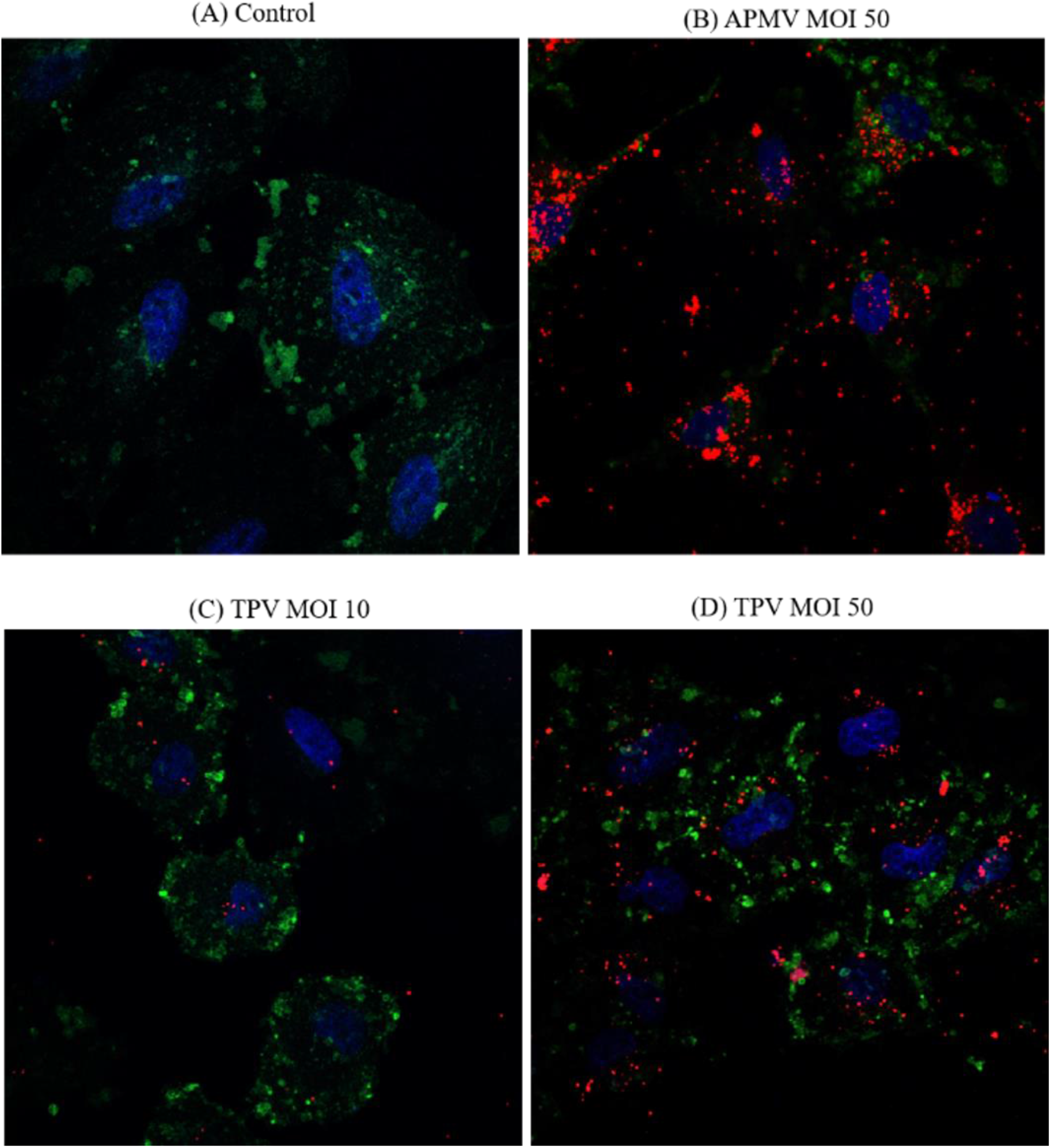
Interaction of APMV and TPV with A549 cells after 36 hours. Fluorescent microscopies showing the A549 cells invasion by APMV and TPV, that were pre-stained with rhodamine-phalloidin (red). A549 cells were incubated with APMV (MOI of 50) and TPV (MOI 10 or 50) for 6 hours. Then the cultures were washed with PBS and filled with DMEM medium for another 36 hours at 37°C. The cells were then washed with PBS and fixed with 4% PFA and labeled with FITC-CTB (green cell membrane) and DAPI (blue-core). The slides were taken to the confocal microscope model TCS-SPE Leica with 63x magnification. **(A)** Cell in DMEM, **(B)** Cells incubated with APMV (MOI 50) presenting a total of 352 dots, **(C)** Cells incubated with TPV (MOI 10) presenting a total of 34 dots and **(D)** Cells incubated with TPV (MOI 50) presenting a total of 447 dots). Scale bar: 10 μm

### TPV induces a small change in cell viability

In view of the results obtained in the confocal fluorescence microscopy suggesting that the GVs were able to be internalized by the cells, the possible repercussions of such an event were analyzed. To this end, we evaluated the cell viability after the interaction of A549 cells with GVs. The protocol adopted in the first microscopy experiment was repeated in this experiment, utilizing unlabeled TPV at MOIs of 0.1, 1, 10 and 50. These results are shown in **Figure 2** by the fluorescence microscopy images obtained from the Live/Dead protocol, as described in Material and Methods. This technique is based on the use of Calcein acetoxymethyl ester (Calcein AM), and Ethidium homodimer-1 (EthD-1) to determine live and dead cells in a culture. Calcein AM is capable of penetrating the cell and being metabolized in living cells emitting a green fluorescence. On the other hand, the EthD-1 is only capable of penetrating the cell membrane that has lost its integrity, thus, as a consequence, it reacts to nucleic acids and generates red fluorescence. **Figure 2A** shows the cell nucleus stained with Hoechst, and **Figure 2B** presents the living cells. As suggested in **Figure 2C**, increasing the number of unlabeled TPV viral particles in the culture results in the increase of dead cells (see increased red marking). The negative control group (maintained in the total absence of viral particles) showed virtually no cell death, while the positive control group (cells incubated with 3% triton) lost 100% of their viability. Cells treated with Triton resulted in complete cell death leading to the saturation of fluorescence, resulting in the bright images observed. **Figure 2D** represents the merging of each group of sampled images. In **Figure 2E**, we performed the quantification of dead cells and it appears to corroborate with the observation that exposure to TPV resulted in cell death. Cultures exposed to an MOI of 10 lost ∼30% of viability, increasing to 40%-50%, in higher MOI of 50. Although it is possible that some of the cells loss of viability was due to viral production, as seen in **Figure 2E**, the viral yield would be so low that it was not susceptible to detection. The values obtained from MOI of 0.1 and 1.0 fall in the statistical margin of error. The proportional increase in cell death to the MOI utilized supports that the virus has a toxic effect over the cells. The statistical analysis showed a p≤0,05, indicating statistical significance.

**Figure 2:**
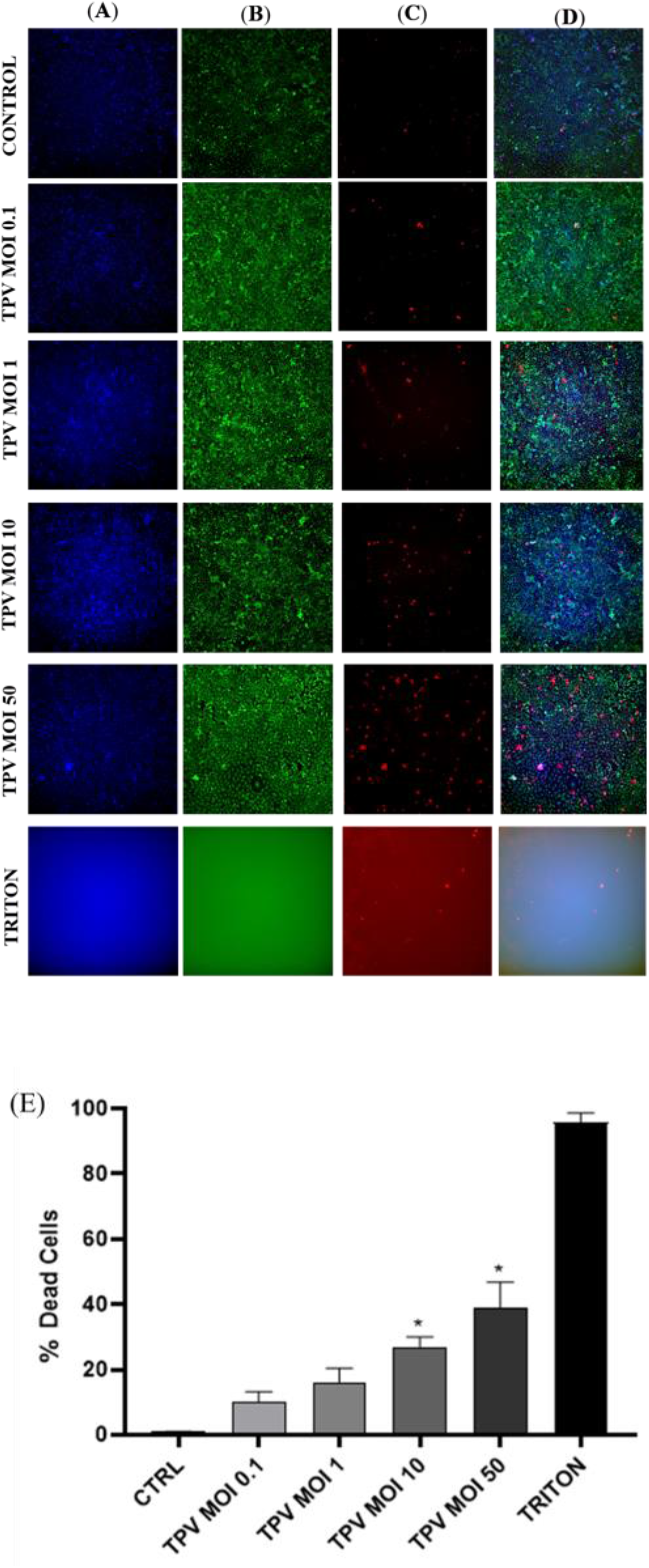
TPV and A549 cell viability. Fluorescent microscopies showing A549 cells incubated with TPV at different MOIs for 6 h and then maintained in DMEM medium. After 36 h, cultures were submitted to the living/dead protocol. Representative images of cell marking. **(A)** DAPI-labeled nucleus, **(B)** green viable cells, **(C)** red dead cells, and **(D)** Merge. Scale bar: 350μM **(E)** Graphic representing the percentage of dead A549 cells related to the presence or absence of TPV, derived from the Live / Dead experiment. All dead cells (red) were multiplied by 100 and then divided by all living cells (green).

### Absence of APMV and TPV replication in A549 cells

As previously noted in the viability experiments with TPV, modulations of immune response pathways or protein synthesis in A549 cells resulting from viral replication steps mostly imply massive loss of viability due to viral replication. Since A549 cells are susceptible to APMV and TPV (**Figure 1**) and that in cell viability experiments (**Figure 2**) dead cells were found in treatments with MOI 10 and 50 of TPV after 36 h, we considered the possibility of this lineage being permissive to viral replicative cycle. Therefore, we kept the viruses in A549 culture for 6 hours at an MOI of 10 or 50 as described previously. After 72 hours of infection, we observed that the morphology of cells exposed to the virus was similar to the control (data not shown), i.e., no cytopathic effect was detected. In comparison, after 72 hours of GV infection in acanthamoeba, ∼100% of cells are lysed (data not shown). Then, supernatants were collected and the adhered cells were lysed. The infected culture material was collected (supernatants and “infected” cell lysates) and subjected to the virus purification protocol. We could not recover viruses from these treatments, as they did not show viral particles after being subjected to centrifugation in a 22% sucrose cushion. Thus, the absence of virions after chemically-induced cell lysis strongly indicates that viral replication is not occurring in cells at the observed time. The lack of viral progeny together with the fact that no notable morphological changes happen during and after viral incubation suggests that, although the cells are susceptible to GVs invasion, they aren’t permissible to replication. However, we cannot rule out that the yield of a possible progeny may exist at a very low titer. Therefore, we do intend to do further experiments to confirm or refute this observation, such as GV genome amplification in “infected” A549 cultures and exposition of such samples to an indicator culture.

### A549 cells are responsive to APMV and TPV fibrils

Since no TPV replication in A549 cells was detected in previous experiments, in order to understand the decrease in cell viability after exposing it to GV, we sought to assess whether APMV or TPV would be able to activate TLR4 receptors on A549 cells. Our rationale assumed in our model that since the virus was seemingly not multiplying in the cell’s interior, replication could not have been the cause of the loss of cell viability observed; and even if there was some residual virus replication, some other/additional factor was probably behind the cytotoxic effect. Due to the similarities between fibril and LPS composition, we theorized that GV-A459 interaction could trigger Toll-Like Receptor 4 (TLR4) cell response. As such, we verified TLR4 expression in A549 cells exposed to APMV (MOI 50) or TPV (MOI 10 or 50) through immunoblotting assays (**Figure 3)**. Bands corresponding to TLR4 were detected in A459-LPS, A459-TPV, and A459-APMV samples (**Figure 3A**). In this experiment, we also used polymyxin B, a cationic antimicrobial polypeptide that inactivates LPS by binding to the lipid A portion of the antigen, inhibiting its activity. We treated the cultures for one hour with 50 μg of polymyxin B and then added TPV (MOI 10 or 50), APMV (MOI 50) or LPS (100 ng/ml) for 30 minutes. As hypothesized, antibiotic treatment protected cells from the effect of LPS and from APMV or TPV exposure, as indicated by the lack of TLR4 recognition-related bands in the immunoblot of treated cells. These results suggest that surface structures of these viruses that are rich in saccharides, i.e., the fibrils are likely to play an important role in the interaction of viruses with the TLR4 (**Figure 3B**). The labelling of the GAPDH was used as loading control for the immunoblot (**Figure 3C**). **Figure 3D** represents the quantification of the antibody TLR4-labeling bands. The experiment showed an increase in the expression of TLR4 protein in samples incubated with the viruses or LPS compared to the control cells. In samples exposed to the viruses, the TLR4 band intensity increase was of ∼2.2x for APMV, and ∼2.3x or ∼3.8x for TPV at MOI 10 or 50, respectively, while samples exposed to LPS increased ∼4.5x. On the other hand, in samples pre-treated with polymyxin B, TLR4 expression was impeded, demonstrated by the lack of bands in the gel. All values were normalized to the GAPDH control. As it can be seen, band intensity increased proportionally to the increase in MOI, with high MOI 50 of TPV presenting an intensity close to pure LPS (3.8 vs 4.5, respectively). These observations argue in favor of the similarity in composition between the viruses fibrils and LPS. All results were statistically significant with p ≤ 0,05.

**Figure 3:**
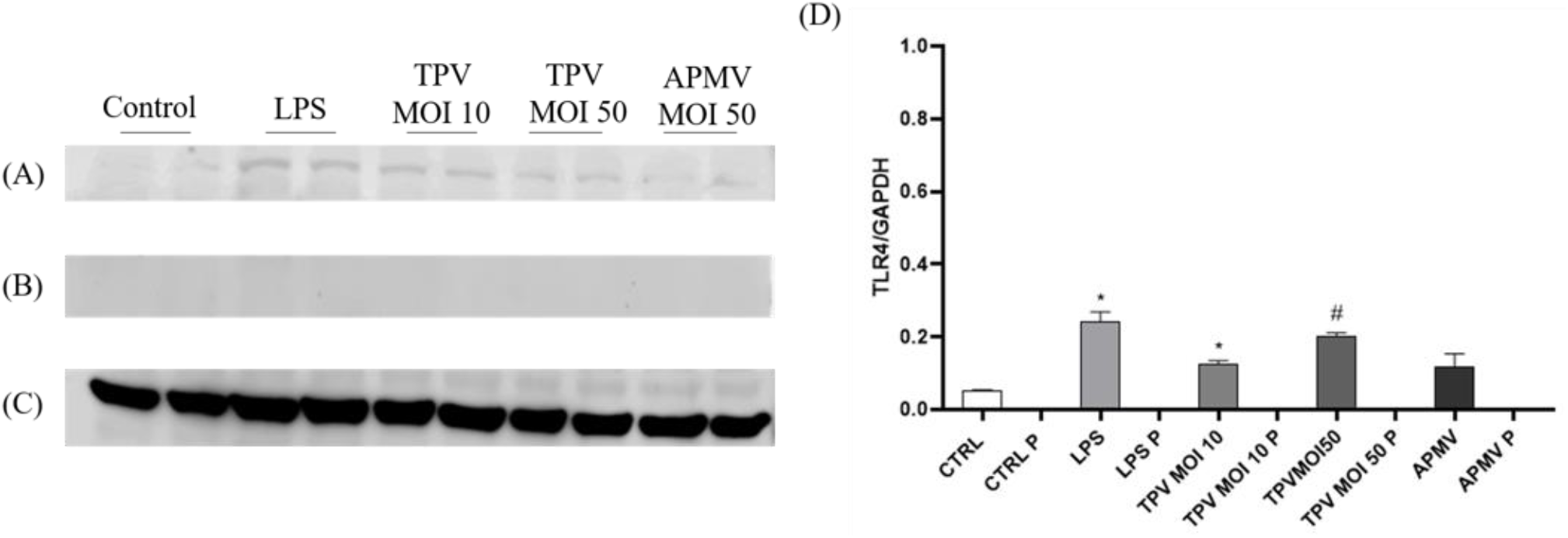
Pulmonary epithelial cells TLR4 are weakly recruited by the presence of APMV and TPV, regardless of the MOI used. Polymixin B completely blocks the activation of TLR4. **(A)** Representative immunoblot membrane that shows the weak TLR4 protein expression in A549 cells exposed for 30 minutes to DMEN, APMV (MOI 50), TPV (MOI 10 or 50), LPS (100 ng/ml). **(B)** Representative immunoblot membrane demonstrating weak expression of TLR4 protein in A549 cells pretreated for 1 hour with polymyxin B (50 μg/m). Then the cells were exposed for 30 minutes to DMEM, APMV (MOI 50), TPV (MOI 10 or 50), LPS (100 ng/ml). All membranes were incubated with anti-TLR4 antibodies. **(C)** To confirm a similar gel load, the membranes were subjected to an anti-GAPDH antibody for two hours. **(D)** Graphic created using the ImageJ processing software the quantification levels of protein expression were made. The data was evaluated as a ratio of the target proteins and the relative expression of the control protein. Student T test was used to determine the statistical significance comparing it with the control: p ≤0,05 *, p ≤0,05 #.

As TLR4 activation can trigger a signaling cascade that ends in IκBα phosphorylation and degradation (Perkins, 2007), we decided to test whether the addition of APMV and TPV are able to activate this pathway. After exposure to the viruses, the pathway activation was verified by the degradation of IκBα protein through western blot, confirmed by the disappearance of the protein band signal as can be seen in **Figure 4**. For cell cultures maintained with TPV at MOI 10 or APMV at MOI 50, the labeling for IκBα was observed, indicating that there was little/no degradation of this protein. However, for cells that received TPV with MOI 50, the labeling pattern was similar to that generated in response to LPS, suggesting greater stimulation of TLR4 and activation of the pathway (**Figure 4A**). A profile similar to that described for TLR4 was observed for cells previously treated with polymyxin, where IκBα degradation seems to have been decreased for the different viruses and LPS (**Figure 4B**). This can be seen by the presence of the IκBα immunoblot band with a greater intensity when compared to the untreated samples, especially in the bands referring to LPS and TPV MOI 50. The GAPDH protein was used as a loading control (**Figure 4C**). **Figure 4D** represents the quantification of the antibody IκBα labeling-band area (Figures 4A and B), with the graphic plotted similarly to the one in **Figure 3D**, but quantifying the loss of band intensity when compared to cells exposed to DMEM. For samples untreated with Poli-B, but exposed to the viruses or LPS the decrease in intensity was of ∼0.42x and ∼0.94x for TPV (MOI 10 and 50 respectively), ∼0.49x for APMV (MOI 50), and ∼0.89x for LPS. While samples treated with polymyxin B showed a decrease of 0.17x and 0.2x for TPV (MOI 10 and 50 respectively), 0.17x for APMV (MOI50), and 0.32x for LPS. It reinforces our observations of change in IκBα presence between untreated and treated samples, as can be seen due to the differences in values obtained.

**Figure 4:**
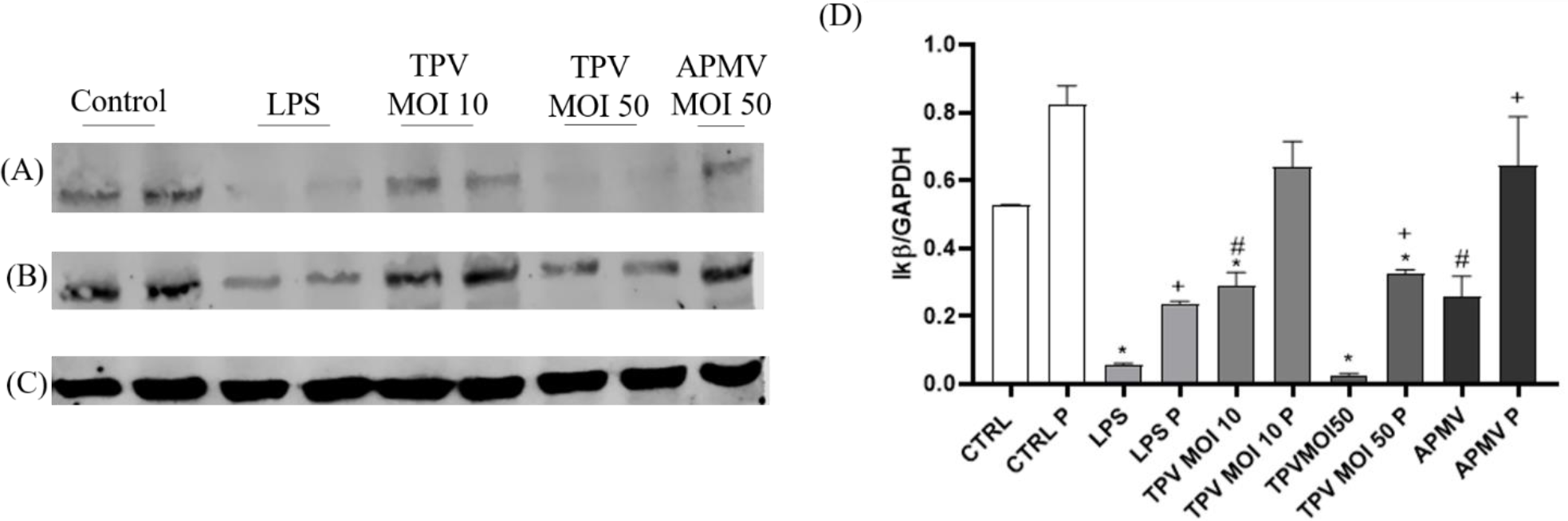
The stimulation with APMV and TPV favors the degradation of IĸBα, while the treatment of A549 cells with polymyxin B decreases the degradation. **(A)** Representative immunoblot membrane demonstrating the absence of IĸBα protein expression in A549 cells exposed for 30 minutes to DMEN, APMV (MOI 50), TPV (MOI 10 or 50), LPS (100 ng/ml). **(B)** Representative immunoblot membrane demonstrating IĸBα protein expression in A549 cells pretreated for 1 hour with polymyxin B (50 μg/m). Then the cells were exposed for 30 minutes to DMEM, APMV (MOI 50), TPV (MOI 10 or 50), LPS (100 ng/ml). All membranes were incubated with anti-TLR4 antibody. **(C)** To confirm a similar gel load, the membranes were subjected to an anti-GAPDH antibody for two hours. **(D)** Graphic created using the ImageJ processing software the quantification levels of protein expression made. The data was evaluated as a ratio of the target proteins and the relative expression of the control protein. Student T test was used to determine the statistical significance comparing it with the control: p ≤0,05 *, p ≤0,05 #, p ≤0,05 +

## 4 Discussion

In this study, we demonstrate for the first time that human type II pneumocytes (A549) can establish adsorptive interactions with members of two genera of GVs: APMV and TPV, resulting in an immunological response. It is important to highlight that epithelial cells act strongly in the maintenance of systemic homeostasis, whether by phagocytizing dead cells or pathogens (Günther and Seyfert, 2018). As such, they are important agents in the innate immune response with the mechanics by which they capture the pathogen being an important factor of their efficacy (Arandjelovic & Ravichandran, 2015). In this context, Chang and collaborators (2020) performed comparative assays of internalization of the bacterium *Klebsiella pneumoniae* (Kp) between macrophages (RAW 264.7) and A549 cells. Their data makes it clear that although epithelial cells can internalize Kp, the process is slow and continuous and therefore dependent only on the time of contact with the pathogen. Macrophages, on the other hand, internalized Kp in less time in a limited way, that is, although bacteria were added to the medium, the internalization rate underwent little change (Chang et al., 2020). This difference can be explained by the absence of specific receptors aimed at phagocytosis in non-professional phagocytic cells, the process being entirely dependent on the pathogen’s virulence factors (Arandjelovic and Ravichandran, 2015; Günther and Seyfert, 2018; Chang et al., 2020). These aspects of phagocytosis in non-professional phagocytic cells are reflected in our results. Here, we observe the internalization of the APMV and TPV viruses (**Figure 1B, C and D**) by A549 cells and we suggest that this internalization may have occurred by phagocytosis, based on information available in the literature (García-Pérez et al., 2003; Arandjelovic & Ravichandran, 2015). In our experiments, we allowed the virus to be in contact with the cells for six hours, and we observed a low uptake in samples that received TPV with MOI 10 (**Figure 1C**) when compared to cells incubated with APMV or TPV with MOI 50 (**Figure 1B, D**). As the process of internalization of pathogens by epithelial cells is slow and continuous (Chang et al., 2020), we believe that a greater number of particles can be internalized with a longer incubation time, even using viruses with a lower MOI. Another interesting observation is the fact that as the MOI increased so did the number of internalized particles, evidenced by the increase of labeled virus signal in the confocal microscopy images (**Figure 1B, D**). This might support the thesis of internalization mediated by phagocytosis due to the high number of particles in these titers not saturating viral entry, which is in accordance with the A549 cells internalization behaviour previously described (Chang et al., 2020). This indicates that it is likely not one specialized receptor behind the capture, but that different non-specific receptors are acting in the process. However, it is necessary to perform further investigation to more consistently support such hypothesis.

After the cell’s invasion, the next step in the replication cycle of viruses is the exposure of their genome to transcription and translation machinery capable of acting on it. During the infection cycle of GV in amoebas, 8 hours after viral adsorption pseudo-organelles called viral factories are observed (Abrahão et al., 2018; Schrad et al., 2020). This region concentrates all the factors necessary for the replication and packaging of the genome during the assembly of viral particles. For APMV and TPV, the release of viral progeny invariably results in cell lysis (Colson et al., 2017a; Oliveira et al., 2019b; dos Santos Oliveira et al., 2021). Also, after ∼4 hours of APMV infection, acanthamoeba cells show a round morphology with loss in adherence as a result of the cytopathic effect promoted by the viral infection. In TPV infection, the cytopathic effect has an additional characteristic, where round cells recruit those not yet infected to form clusters known as grape bundles (Oliveira et al., 2019b). In A549 cells, no morphological changes were observed during the 42 hours of infection. In addition, the DAPI used to label cell nuclei in confocal microscopy experiments is also capable of marking viral factories. However, our experiments did not reveal this additional labelling, suggesting that viral factories are absent in cells that have internalized the virus (**Figure 1**). When put together, this information supports the failure to obtain viral particles in A549 cells incubated with APMV or TPV. It is also possible that, if any replication occurs in the interior of A459 cells, the dynamics of infection may be different and we were not able to detect the viral factory in the images acquired. Thus, a kinetics of infection will be performed in a subsequent future work.

There is strong evidence supporting the previous observation of the epithelial cell line used, seeming to be susceptible, but not permissive, to APMV and TPV. There are a number of possible reasons for the lack of replication, with one of them being that the epithelial cell did not possess the appropriate cellular factors for the viral genome replication. Another possible explanation is that the degradation of the virus particle by A459 occurs before its infective cycle is completed, thus replication is inhibited (Lambotin et al., 2010). It is unlikely that the GVs particles would be easily degraded due to its notorious particle resistance (Sharma et al., 2020), with further investigations being necessary to determine the possible fate of internalized viruses. One plausible course of action would be the amplification of viral genomes from infected A459 cultures, as mentioned previously.

After concluding based on our evidence that replication was not occurring, we sought information about the physiology of the A549 cells in order to understand the 40-50% reduction in viability caused by the interaction with TPV in the highest MOI used (**Figure 2A, B**). The literature has shown that A549 cells have both TLR4 and MD-2 mRNA expression (Guillot et al., 2004; Sender & Stamme, 2014). Strong inflammatory response mediated by TLR4 is associated with cellular toxicity (Olejnik et al., 2018), therefore it is a plausible explanation for the observed cellular death. Generally speaking, the lipopolysaccharide binding protein (LBP) binds and transfers LPS to CD14. The CD14 protein has the function of transferring LPS to the MD-2 protein (a glycoprotein that is associated with TLR4). The MD-2/LPS interaction causes a structural change in the TLR4/MD-2 complex, which allows the release of NF-kB via the MyD88-dependent pathway (Lu et al., 2008; Thorley et al., 2011). We know that APMV fibrils have carbohydrates in their composition and some of these are also found in bacterial surface structures, such as LPS. Examples of such shared carbohydrates include rhamnosamine and viosamine (Piacente et al., 2012; Parakkottil Chothi et al., 2010). Similar to APMV, TPV also has fibrils on its surfaces that in turn also have polysaccharides in their composition, although, different from APMV, their composition has yet to be determined. Therefore, our hypothesis is that these surface structures of APMV and TPV, present structural similarities that are capable of triggering TLR4 activation, most likely by mechanisms similar to those of bacteria. This hypothesis is corroborated by our observation that both the addition of bacterial LPS and the maintenance of the virus in culture for 30 minutes timidly initiate the increase in TLR4 expression (**Figure 3A, D**). These results support not only the hypothesis that they activate the TLR4 signaling pathway, but also further poses the notion that both viruses’ fibrils possibly share some structural similarity to bacterial LPS. The small increase in TLR4 expression in the presence of LPS or viruses may be related to the low availability of CD14 in this cell line. Previous studies reported that A549 cells, unlike macrophages, have little expression of membrane CD14, requiring the addition of this protein to the culture through the supplementation of fetal bovine serum (FBS) (MacRedmond et al., 2005). Future studies can make a deeper analysis of the participation of these proteins in TLR4 activation necessary. On the other hand, cells treated with the LPS inhibitor (Polymyxin B) were protected from the action of LPS and viruses, since, in this case, there was no increase in TLR4 protein expression (**Figure 3B, D**). The results obtained for the IκBα protein points in the same direction as the previous experiment. The IκBα degradation pathway, activated by the interaction of LPS with TLR4, also seems to be stimulated by TPV (MOI 50) and, to a lesser extent, from exposure to TPV (MOI 10) and APMV (MOI 50). For this situation, there is a reflection similar to the previous one: the low activation of TLR4 can occur due to the low expression of CD14 and this directly interferes with the result (**Figure 4A, D**). As was observed for TLR4, pretreatment with Polymyxin B decreased IkBa degradation (**Figure 4B, D**).

Besides the epithelial cell interaction, Abrahão and collaborators (2018) had already reported that TPV establishes cytotoxic interactions with *Tetrahymena* sp and acanthamoeba (Abrahão et al., 2018). Classified as unicellular ciliated protozoan, the *Tetrahymenas* inhabit fresh water, have between 30 to 50 μm in length and are considered professional phagocytic organisms (Ruehle et al., 2016). In their studies, Abrahão et al. have demonstrated that similar to what was observed for A549 cells, the protozoan cells are capable of capturing TPV particles, but are not capable of replicating its genome. From an elegant approach, it was made clear that after adsorption, the ribosomes are separated from the cytoplasm through the formation of a large vesicular area, a product from the agglomeration of numerous vesicles initially present around the nucleus. Simultaneously, the nucleus is also a target of degradation which would help the eventual separation of ribosomes. The ribosomal RNA degradation was denominated ribosomal shutdown (Abrahão et al., 2008; Rolland et al., 2020). In our work, a similar cytotoxicity effect due to exposition to the viruses was witnessed for A549 cells as exposure to both APMV and TPV, even in the apparent absence of replication. However, this phenomenon cannot be attributed to a ribosomal shutdown due to the lack of evidence, probably being a result of the triggering of inflammatory response via TLR4 mediated pathway activation. The lack of replication indicates that such effects are consequential to some structural aspect inherent to the virion, quite possibly the fibrils.

In conclusion, this work has obtained some intriguing data that reinforces the peculiarity of the interaction between GVs and human cells, as well as of these viruses structures. Taken our data together, we suggest that upon interaction with mimiviruses, mammalian cells recognize viral structures, probably the fibrils, triggering a TLR mediated MyD88 signalling cascade. This pathway leads to the IκBα degradation (**Figure 4 A, E**) and release of NFkB to the nucleus, resulting in the expression of proinflammatory cytokines as summarized in **Figure 5**. Previous genomic and proteomic studies based on data obtained from genome sequencing show that mimiviruses possess genes related to organisms present in each of the three domains of life. This amazing aspect reinforces the possible evolutionary relationships of the GVs with bacterias, archeas, and eukaryotes. Such data can help enlighten the similarities observed in structural composition, and how these structures can be related with infection mechanisms or the activation of cytotoxic response in different cell types (Raoult et al., 2010; Abrahão et al., 2018; Miranda Boratto et al., 2019). The results shown here seem to reinforce this possible evolutionary relationship, as both APMV and TPV appear to trigger the TLR4 response similarly to the LPS, most likely due to structural similarities. Our results serve as a base to further explore this very complex relationship.

**Figure 5:**
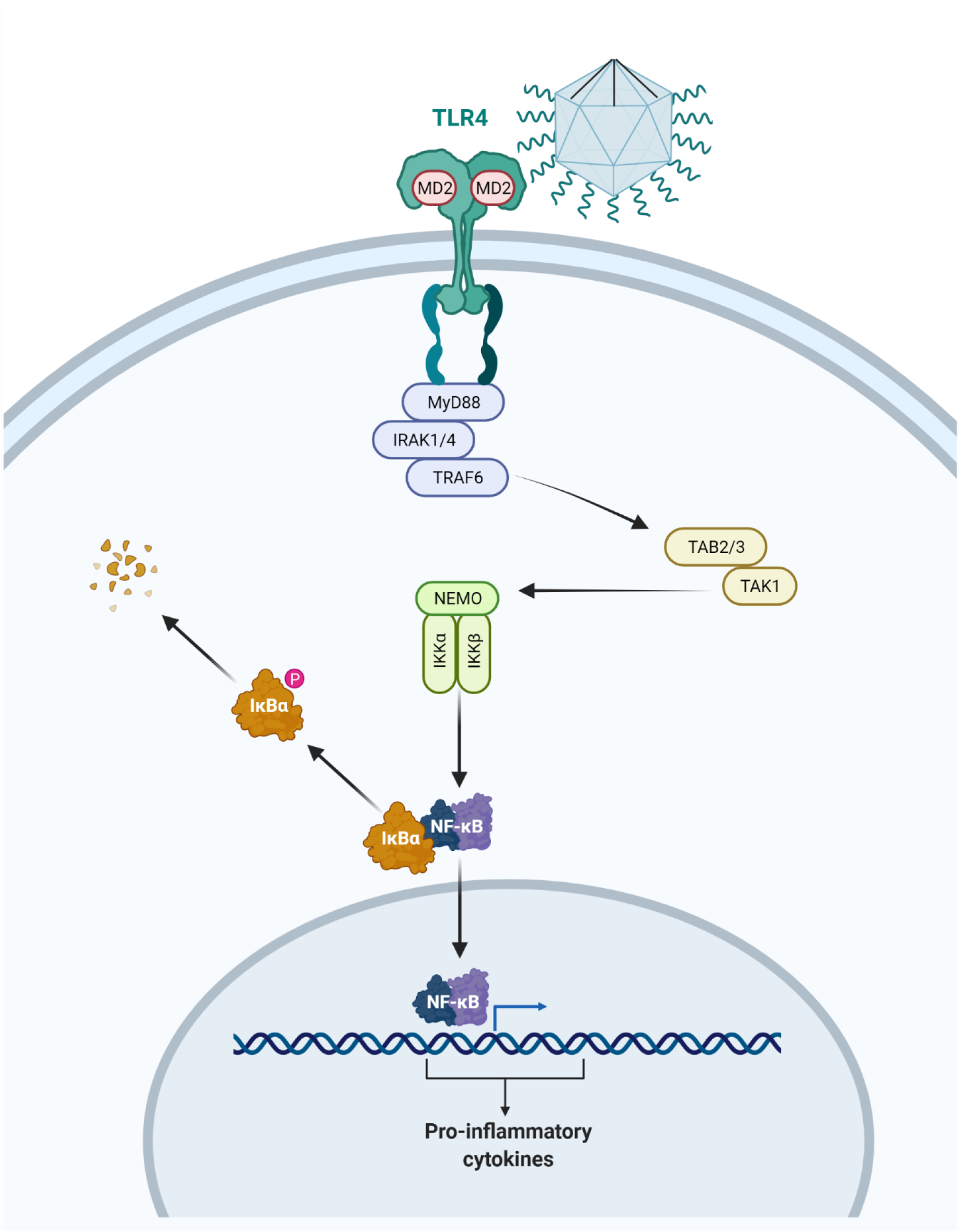
Illustration representing the activation of the myeloid 88 dependent TLR4 pathway by contact with the APMV. Following the TLR4 activation, the IRAK4 kinase suffers phosphorylation and activates IRAK1 resulting in TRAF6 recruitment. This allows the TAK1 and TAB2/3 complex to activate the kinases IKKα, IKKβ, and IKKγ. These enzymes promote the IkB phosphorylation and its degradation allows the translocation of the nuclear factor-kB (NF-kB) to the nucleus, and the production of inflammatory cytokine. This representation is merely illustrative and is not up to scale. Drawing created with BioRender.com.

## 4 Acknowledgments

We thank the National Center for Structural Biology and Bioimaging, in particular the Plataforma de High Content/throughput Analysis, for providing an easily accessible infrastructure for research and for the detailed and insightful analysis of the data.

## 6 Author Contributions

JSO performed experiments and wrote the manuscript. DFO performed experiments. VAE drew the ilustration and revised the text. GHPN revised the text. LO helped with the microscopy. LN, JMP, AJG and DF helped in the development of the experiments.

