## Supplementary Material for "Paving the way towards understanding the inflammatory pathways triggered by giant viruses in mammalian cells: effect of mimivirus-cell interactions on IκBα degradation"

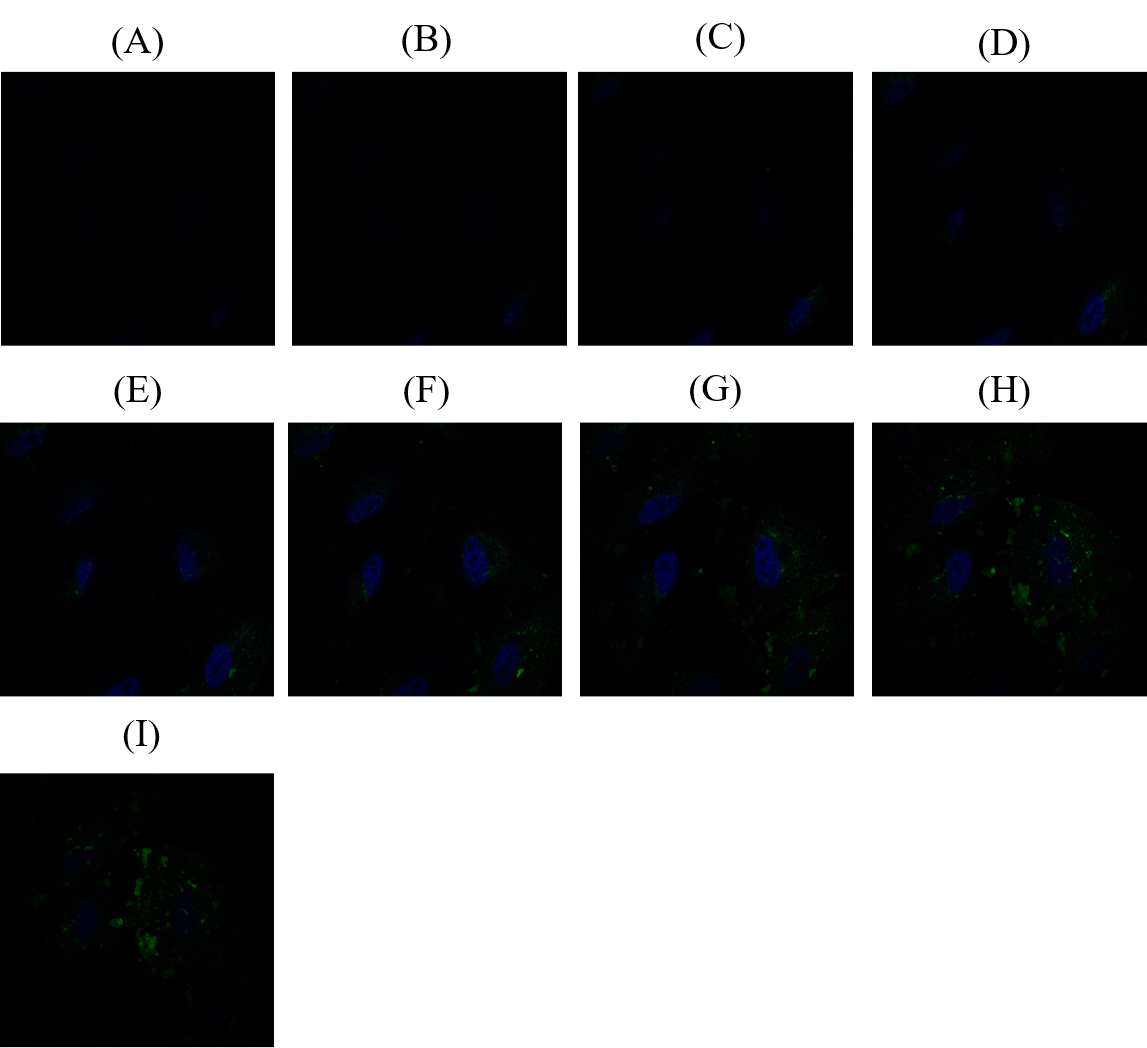

**Supplementary Figure 1.** *A549 cells exposed to DMEM.* 10,000 cells were plated in 24-well plates with coverslips for 24 hours. Subsequently, cultures were washed with PBS and filled with DMEM for 6 hours. Then, cultures were washed again and kept in DMEM for another 36 hours. **(A-I)** Confocal microscopy Z cell images. DAPI was used to label the nuclei (blue fluorescence) and fluorescein isothiocyanate (FITC) -cholera toxin B subunit (CTB) was used to label the cell membrane. Z-stack images were collected to a total of 9 optical slices with an interval of 0.7 μm, at a position from -10.5 μM to -16.1 μM. The images represent the slices positioned at **(A)** -10.5μM; **(B)** -11.2 μM; **(C)** -11.9 μM; **(D)** -12.6 μM; **(E) -**13.3 μM; **(F)** -14.0 μM; **(G)** -14.7 μM; **(H)** -15.4 μM; **(I)** -16.1 μM. Scale bar: 10 μM.

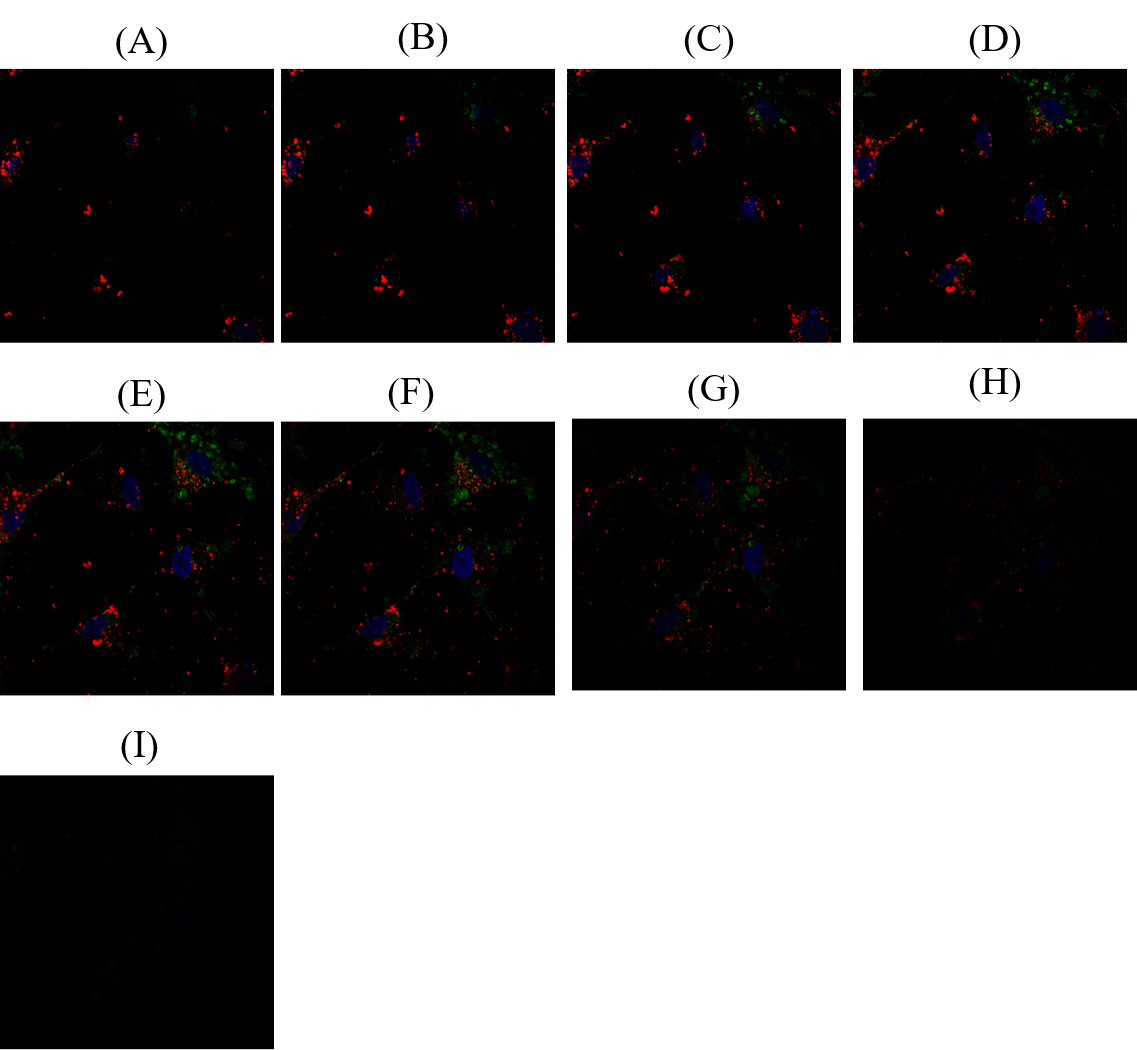

**Supplementary Figure 2.** *A549 cells interact with APMV*. A549 cells interact with TPV (MOI 10). 10,000 cells were plated in 24-well plates with coverslips for 24 hours. Subsequently, cultures were washed with PBS and filled with DMEM containing the APMV particles (MOI of 50) for 6 hours. Then, cultures were washed again and kept only in DMEM for another 36 hours. **(A-I)** Confocal microscopy Z cell images, showing the interaction of viruses with cell membranes. DAPI was used to label the nuclei (blue fluorescence) and fluorescein isothiocyanate (FITC) - cholera toxin B subunit (CTB) was used to label the cell membrane and rhodamine phalloidin used to label the viral particles. Z-stack images were collected to a total of 9 optical slices with an interval of 0.7 μm, at position from -4 μM to -9.6 μM. The images represent the slices positioned at **(A)** - 4 μM; **(B)** - 4.7 μM; **(C)** -5.4 μM; **(D)** - 6.1 μM; **(E) -** 6.8 μM; **(F)** -7.5 μM; **(G)** -8.2 μM; **(H)** -8.9 μM; **(I)** -9.6 μM. Scale bar: 10 μM.

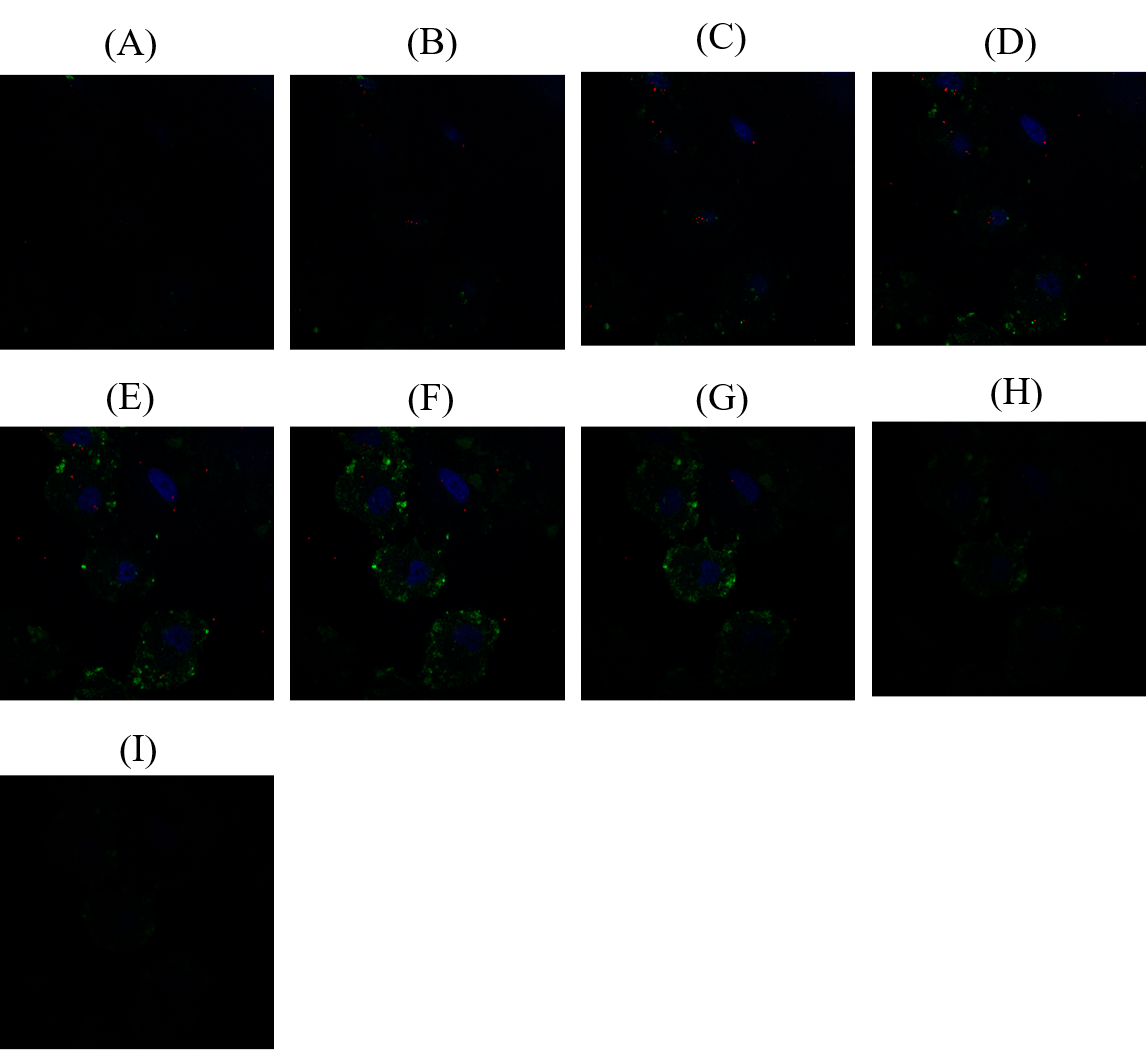

**Supplementary Figure 3.** *A549 cells interact with TPV (MOI of 10)*. 10,000 cells were plated in 24-well plates with coverslips for 24 hours. Subsequently, cultures were washed with PBS and filled with DMEM containing the TPV particles for 6 hours. Then, cultures were washed again and kept only in DMEM for another 36 hours. **(A-I)** Confocal microscopy Z cell images, showing the interaction of viruses with cell membranes. DAPI was used to label the nuclei (blue fluorescence) and fluorescein isothiocyanate (FITC) - cholera toxin B subunit (CTB) was used to label the cell membrane and rhodamine phalloidin used to label the viral particles. Z-stack images were collected to a total of 9 optical slices with an interval of 0.7 μm, at position from -24.2 μM to -29.8 μM. The images represent the slices positioned at **(A)** – 24.2 μM; **(B)** – 24.9 μM; **(C)** -25.6 μM; **(D)** – 26.3 μM; **(E) -** 27 μM; **(F)** - 27.7 μM; **(G)** -28.4μM; **(H)** -29.1 μM; **(I)** -29.8 μM. Scale bar: 10 μM.

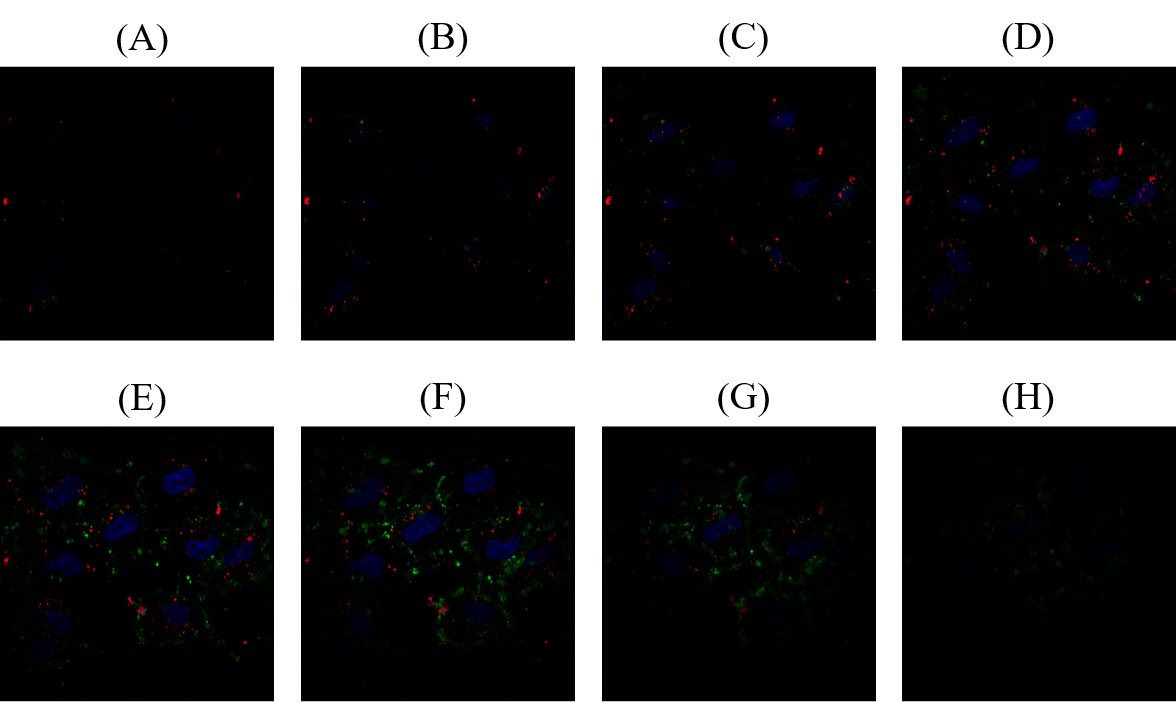

**Supplementary Figure 4.** *A549 cells interact with TPV (MOI of 50).* 10,000 cells were plated in 24-well plates with coverslips for 24 hours. Subsequently, cultures were washed with PBS and filled with DMEM containing the TPV particles for 6 hours. Then, cultures were washed again and kept only in DMEM for another 36 hours. **(A-H)** Confocal microscopy Z cell images, showing the interaction of viruses with cell membranes. DAPI was used to label the nuclei (blue fluorescence) and fluorescein isothiocyanate (FITC) - cholera toxin B subunit (CTB) was used to label the cell membrane and rhodamine phalloidin used to label the viral particles. Z-stack images were collected to a total of 8 optical slices with an interval of 0.7 μm, at position from 1.3 μM to -3.6 μM. The images represent the slices positioned at (**A)** 1.3 μM; **(B)** 0.6 μM; **(C)** -0.1 μM; **(D)** – 0.8 μM; **(E) -** 1.5 μM; **(F)** – 2.2 μM; **(G)** -2.9 μM; **(H)** -3.6 μM. Scale bar: 10 μM.
